# The ABCD Stop Signal Data: Response to Bissett et al.

**DOI:** 10.1101/2020.07.27.223057

**Authors:** H Garavan, B Chaarani, S Hahn, N Allgaier, A Juliano, DK Yuan, C Orr, R Watts, TD Wager, O Ruiz de Leon, DJ Hagler, A Potter

## Abstract

This paper responds to a recent critique by Bissett and colleagues (Bissett et al., eLife, In Press) of the fMRI Stop task being used in the Adolescent Brain Cognitive Development^SM^ Study (ABCD Study^®^). The critique focuses primarily on a design feature of the task that the authors contend lead to a violation of race model assumptions (i.e., that the Go and Stop processes are fully independent) which are relevant to the calculation of the Stop Signal Reaction Time, a measure of the inhibition process. Bissett and colleagues also raise a number of secondary concerns. In this response we note that satisfying race model assumptions is a pernicious challenge for Stop task designs but also that the race model is quite robust against violations of its assumptions. Most importantly, while Bissett et al. raise conceptual concerns with the task we focus here on analyses of both the performance and the neuroimaging data and we conclude that the concerns appear to have minimal impact on the neuroimaging data (the validity of which do not rely on race model assumptions) and have far less of an impact on the performance data than the critique suggests. We note that Bissett et al. did not apply any performance-based exclusions to the data they analyzed, that a number of the trial coding errors that they flagged were already identified and corrected in the ABCD annual data releases, that a number of the secondary concerns reflect sensible design decisions and, indeed, that their own computational modeling of the ABCD Stop task suggests the problems they identify have just a modest impact on the rank ordering of individual differences in subject performance. In this paper, we list some adjustments that have been made to the task and some new flags that are now added to the annual, curated data releases. We stress that the ABCD data are fully available to the scientific community who are empowered to apply whatever inclusion and exclusion criteria they deem appropriate for their analyses and we conclude that the ABCD Stop task yields valuable data that researchers can use to track adolescent neurodevelopment.

---

Bissett and colleagues (Bissett et al., eLife, In Press) list a number of concerns regarding the specific version of the Stop Signal Task that is included as one of three fMRI tasks in the neuroimaging battery of the multi-site, longitudinal Adolescent Brain Cognitive Development (ABCD) study (www.ABCDstudy.org). The ABCD study is committed to full sharing of its tasks, data processing and analysis scripts, and dataset. We are delighted to see “open science” in action, and we value the scientific community flagging concerns and helping us to continually improve this landmark study.

The Bissett et al. critique focuses largely on stimulus design characteristics of the ABCD Stop Signal Task which may lead to violations of race model assumptions (Logan et al., 1984). The concerns are well described in Bissett et al. In brief, the onset of the Stop signal, which is controlled by an adaptive, performance-related algorithm, is accompanied by the simultaneous offset of the Go choice stimulus. One consequence of this design feature is that the on-screen duration of the Go choice stimulus can be shorter on Stop trials. The concern raised by Bissett et al. is whether this might affect the context independence of the Go process, that is, that the Go process is unaffected by whether or not a Stop signal is presented on a trial. Context independence is assumed by race model theories of response inhibition and underpins the valid calculation of the Stop Signal Reaction Time (SSRT), an estimate of the duration of the stopping process. In addition to this primary issue, Bissett and co-authors raise a number of other concerns with the task design and conclude that these “significantly compromise” the value of the data. We disagree with this conclusion and provide evidence in support of the utility and validity of the Stop task data from the ABCD Study. The Bissett et al. critique raises important theoretical issues related to the assumptions of the race model underlying calculation of the SSRT, but the issues raised do not necessarily undermine the utility of the ABCD SSRT estimates as a measure of individual differences. Importantly, Bissett et al. do not examine whether the issues they raised do in fact affect the ABCD study’s SSRT estimates, nor whether they have an impact on the brain imaging data.

While acknowledging the concerns regarding race model violations, we focus here on empirically investigating the extent to which these violations meaningfully impact the quality of the ABCD data. We present a series of analyses of both the SSRT estimates and the validity of the neuroimaging data. Ensuring independence of the Go and Stop processes is a perennial concern with the Stop task and not one peculiar to the ABCD task version as Bissett and colleagues have themselves argued (Bissett et al., 2021). Moreover, the Stop task is quite robust against violations of race model assumptions (Band et al., 2003). Thus, it is critical to determine whether, or to what degree, the concerns raised do indeed corrupt the SSRT estimates. We demonstrate here that they appear to have only modest effects on SSRT and, furthermore, do not substantively impact or invalidate the Stop task fMRI measures.

Some of the concerns raised by Bissett et al. were in fact known issues that were corrected prior to the annual data analyses and every curated data release of the ABCD Data Analysis, Informatics and Resources Center. Bissett et al. applied no performance flags (either their own or those recommended in all ABCD data releases) that serve to exclude participants who do not perform the task appropriately. Other recommended corrections, such as flipping left-right responses in those participants with reversed response paddles, were not applied. And, finally, we contend that many of the minor issues raised by Bissett et al. are design features and not design flaws. Following the format of the original paper, we address each issue in turn and offer our recommendations on if, and how, each might be addressed. The ABCD task fMRI working group in consultation with members of its External Scientific Board and outside experts have decided to make a number of small changes to the task. These changes are in consideration of the impact of the issues raised by Bissett et al. balanced against the implications of changing a task once a longitudinal study has commenced.

## ABCD Stop task

The code underlying the analyses conducted in this paper are available (https://github.com/sahahn/SST_Response). We underscore that although the annual ABCD data releases provide updates and contain flags identifying problematic data (e.g., poor quality images, poor task performance), the data are fully available to the scientific community empowering researchers to apply whatever inclusion and exclusion criteria they deem appropriate for their specific research questions. Figure 1 shows the performance criteria that have been used to calculate the Stop task performance flag that is part of the recommended inclusion criteria for these data provided with each ABCD data release. Figure 1 shows two important properties of the ABCD Stop task data. First, performance statistics show that the tracking algorithm achieves a successful inhibition rate of approximately 50% (which maximizes efficiency of SSRT estimation), few omission errors, and typically distributed response times. Second, it shows robust stopping-related activation in known response inhibition-related regions. These characteristics increase confidence that the ABCD Stop task data offer insightful measures of inhibitory control abilities and related brain function.

**Figure 1:**
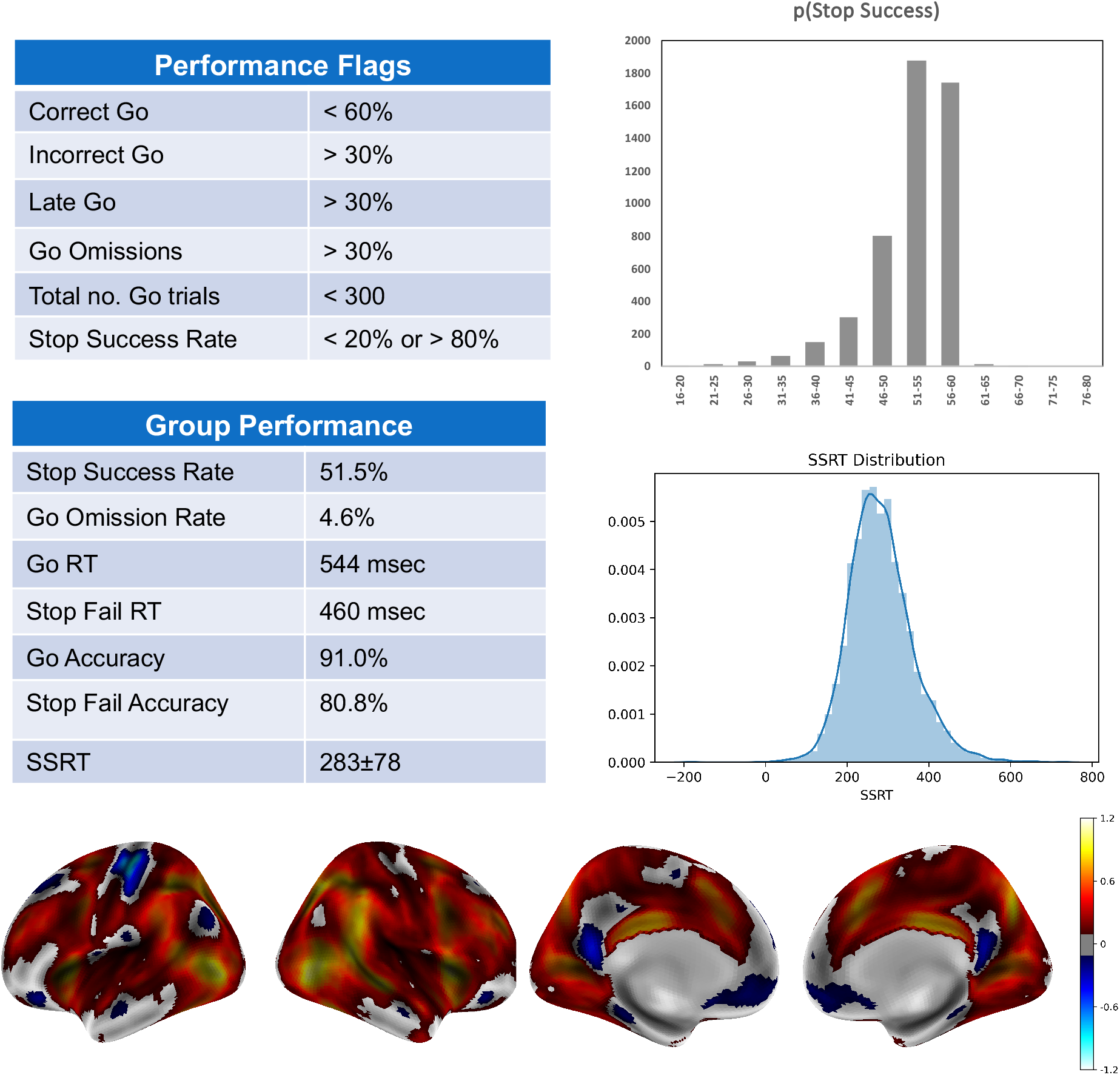
ABCD Stop Signal Task. The performance criteria (recommendations for participant exclusion) and group level performance on the ABCD Stop Signal Task are shown in the tables. Histograms of the p(Successful Stopping) and SSRT and group activation maps (Cohen’s d threshold of .2 for Successful Stops vs Correct Go trials), for the baseline (age 9 and 10) data are shown.

### Issue 1. Different go stimulus duration across trials

As noted above and described in Bissett et al., the offset of the Go choice stimulus coincident with the onset of the Stop signal may reduce the strength of the Go process on Stop trials, violating context independence. Poorer choice accuracy on Stop Fail trials compared to Go trials (Figure 1), especially at shorter Stop Signal Delays, indicates that this is a valid concern; the poorer choice accuracy on the Stop Fail trials is consistent with the Go process being different on these trials in comparison to Go trials. In our analysis, with performance criteria applied and the exclusion of a small number of participants who experienced a task programming error (see Issue 3 below), Go accuracy = 91% and Stop Fail accuracy = 81%.^1^

The critical issues are to what extent this violation of race model assumptions impacts reaction time and/or brain imaging data, and whether any violation impacts the utility of SSRT as a measure of individual differences in inhibitory control. Full context independence is difficult to attain. Bissett et al. and others demonstrate that this is the case across many Stop Signal Tasks, including those without the stimulus design feature of the ABCD task (Gulberti et al., 2014; Bissett et al., 2021). The presentation of a Stop signal has the potential to impact an ongoing Go response, even if the Go stimulus remains on screen. In addition, it is known that participants slow their Go responses in anticipation of a Stop trial (indeed, this “proactive” control can be modelled; Harlé et al., 2016). “Strategic” adjustments in the speed of Go trial responding can vary within an experiment and across individuals, and can affect SSRT estimates (Leotti et al., 2010).

One test for clear violations of context independence, which Bissett et al. presents, is whether Stop Fail response times (RTs) are slower than Go RTs. This pattern cannot be explained by the standard race model, which assumes Go processes are equivalent on Go and Stop signal trials and, further, that the observed distribution of RTs on Stop Fail trials are censored (by virtue of the Stop process completing before the relatively slower Go responses would be made). As Bissett et al. note, the ABCD data pass this test: Stop Fail trials are not slower than Go response times. Table 1 shows RT data for ABCD as calculated by Bissett et al., after we applied the standard performance flags, and excluded participants affected by the Issue 3 programming error (described below). Stop Fail RT is 83 msec faster than Go RT (t_(1,7114)_ = 53, p < .0001), broadly consistent with context independence. For comparison, we include two datasets from Bissett and colleagues, chosen because they do not share the ABCD design property of offset of the Go stimulus with the onset of the Stop stimulus. We refer to these datasets as the Ontology study (n = 522; Eisenberg et al., 2019) and the Phenome study (n = 130, healthy controls only; Poldrack et al., 2016). We use the low frequency condition (20% Stop trials) in the Ontology study as it is closer to the ABCD proportion of 17%. The ABCD value of 83 msec is within the range of these other studies: Stop Fail is 30 msec and 122 msec faster than Go trials in the Ontology and Phenome studies, respectively.

**Table 1:** Performance statistics relevant to Issue 1 for ABCD and two comparison datasets (see text for details). We replicate the initial calculations by Bissett and colleagues of the mean response times for Go RT and Stop Fail (SF) RT trials, we apply the exclusion of participants based on poor performance as recommended with the ABCD annual data releases, and then we drop the RT data of an additional 1.24% of participants whose task was compromised by a coding error (described under Issue 3 below).

|  | ABCD |  |  | Comparison |  |
| --- | --- | --- | --- | --- | --- |
|  | Initial | Exclude poor performers | Issue 3 exclusion | Ontology | Phenome |
| Mean Go RT | 543 | 543 | 544 | 571 | 478 |
| Mean SF RT | 459 | 458 | 460 | 541 | 356 |
| Difference | 84 | 85 | 83 | 30 | 122 |

<sup>1</sup> Excluding participants who fail to adequately comply with task instructions or who show aberrant performance is a standard practice and especially important given the young age of ABCD participants. Recommended performance criteria that accompany ABCD data releases are shown in Figure 1 but these were not applied by Bissett et al. In addition, a programming error detailed under Issue 3 led to a very different task experience for certain participants and we are recommending that these participants (1.24% of the sample) be excluded from analyses.

Next, Bissett et al. reports that 6.2% of the ABCD participants do not show the expected RT pattern and label them as violators (i.e., participants for whom Stop Fail RT > Go RT). While one might refine this estimate (it drops to 5% once performance flagged participants and participants on whom there was a task programming error described below are excluded) it is nonetheless superior or comparable to the estimates for the Ontology and Phenome studies (17% and 4.4%, respectively). These percentages may not indicate statistically reliable effects (i.e., the differences in RTs may not differ reliably from 0), as confidence intervals and estimates of their sampling distribution are not provided. Although analyses of individual participants are likely underpowered, with too few trials for robust behavioral analyses, one can estimate the numbers of participants who might be deemed true violators (i.e., with a significant one-tailed t-test per participant comparing Stop Fail trial RTs against Go trial RTs). The percentage is low: 1.6% for ABCD, again falling between the Ontology (2.7%) and Phenome (0%) studies that were designed without coincident Go stimulus offset and Stop stimulus onset.

We conclude that some number of participants evidencing context independence violations are to be expected in many Stop task designs. While researchers can make their own decisions on participant inclusion and exclusion criteria given ABCD’s data sharing procedures, we note that a recent Stop task “best practices” paper recommends that SSRT should not be estimated for those participants who violate the Stop Fail RT < Go RT criterion (Verbruggen et al, 2019). Consequently, a new flag identifying the 5% of ABCD participant “violators” is included in the annual ABCD data releases (see Implications and Recommendations below). Note that this flag is applied to any participant whose Stop Fall RT > Go RT by any amount (i.e., 1 ms or greater).

#### Individual Differences

Examining individual differences is a central goal of the ABCD study. Crucially, for the utility of the ABCD SSRT estimate to be degraded as a measure of individual differences, violations of context independence must result in more than a shift in mean SSRT. Rather, the rank ordering of participants’ SSRT values must be substantively altered. As all ABCD participants performed the same task, we expect individual differences to be largely unaffected by the abbreviation of the Go process as a function of task structure, but this expectation awaits further experimental or computational modeling studies. Computational modeling approaches have potential to characterize the specific design features of the ABCD Stop task and the additional processes that can occur on all Stop tasks (e.g., “trigger errors”, or trials on which the STOP process is never initiated; Weigard et al., 2019). Bissett et al describe some preliminary drift diffusion models to capture the particulars of the ABCD task design. While we leave it to others to judge the value of these models, we note that they return “adjusted” SSRT estimates that don’t, in fact, appear to be that different to the estimates derived from the standard analyses that assumes context independence. To elaborate, an essential element of the Bissett et al model is the estimate of inter-individual variation in SSRT. To assign an SSRT value to their simulated subjects, they “sampled randomly from an SSRT distribution with a mean that equaled the observed ABCD grand mean but assumed four different levels of between-subject variability (ranging from SD = 0-85ms).” Their own analyses of “20 simple stopping conditions from a recent large-scale stopping study” estimated the mean between-subject SD of SSRT to be 43 msec with a range of 28-85 msec. At the higher estimate (85 msec) they estimate the mean rank correlation between SSRT as calculated by the Independent Race Model (i.e., assuming context independence) and their three alternative models to be .93. Unfortunately, they did not report the correlation for 43 msec although this is the empirical mean that they estimated from their own analyses of their 20 simple stopping conditions. The minimal estimate from their analyses is 28 msec but the correlation for this value is also not reported. Instead, the nearest estimate to the bottom of their observed range (25 msec) yields a mean correlation of .78. The other estimates that they report in their paper, and to which they pay specific attention, are 5 msec and 0 msec. However, these are far beyond the range of their empirically observed estimates and are not at all credible estimates of the true inter-subject variability in SSRT. Using their code, we have repeated their analyses using their mean estimate of SSRT SD (43 msec). We observe a mean correlation between their computational models and the ideal independence race model to be .85. The true estimate of inter-subject variability in SSRT in the ABCD data is, of course, uncertain if one holds that these SSRT estimates are invalid. Nonetheless, if we exclude violators, remove participants who experienced a task programming error (see Issue 3 below), exclude participants flagged for poor performance, and calculate SSRT with the 0 msec SSD trials excluded (see Issue 2 below), we calculate the SSRT SD to be 73 msec. This estimate, in turn, yields a mean correlation between their computational models and the ideal independence race model to be .91. Thus, from the simulations and computational modeling conducted by Bissett et al, we conclude that modeling the context violation present in the ABCD study is unlikely to distort the rank ordering of participants in a meaningful way.

Future experimental or computational modeling studies could help provide greater clarity about these matters. Until that time, researchers are encouraged to carefully consider the assumptions of any measurement model they apply to these data, including the race model-based SSRT estimate, and to consider the possible limitations of parameter estimates and measures derived from any model that assumes context independence.

#### Stop Task Brain Activation

Turning to the brain activation data, it is important to note that the measurement assumptions that the activation contrasts reflect valid measures of response inhibition are much simpler than those required for the SSRT estimation and do not rely on race model assumptions, including context independence. Brain activity can be associated with response inhibition processes if one compares trials requiring inhibition of prepotent responses against trials that do not. A standard contrast to achieve this end for ABCD would be to compare Successful Stop trials against Go trials. The shorter duration of Go choice stimuli on Stop trials compared to Go trials does introduce differences between the two conditions. However, there are typically other more substantial differences present when isolating inhibition-related activation: There is a motor response on Go trials and not on Stop success trials, only the latter contains a Stop signal, and so on. The contrast of Successful Stop trials against the implicit baseline and the contrast of Successful Stop trials against Failed Stop trials are also available to researchers. As is always the case, researchers using these data should be aware of design specifics and determine if they impact on the researcher’s specific question. The ABCD Stop task has already been shown to produce robust activation in the response inhibition network and activation levels show the anticipated correlations with individual differences in SSRT (Casey et al., 2018; Chaarani et al., In Press). A very similar task, with the same Go stimulus design features, has been employed in the IMAGEN study of adolescent development (Schumann et al., 2010) and has, for example, identified functional differences between adolescents with substance use, adolescents with ADHD, adolescents with psychotic symptoms, dysregulated youth and controls (Bourque et al., 2017; Specher et al., 2019a; Whelan et al., 2012), and has predicted future drug use (Spechler et al., 2019b; Whelan et al., 2014).

We demonstrate the validity of the brain activation measures in ABCD with two analyses. The first examined brain activation in the violators identified above (a “worst case” scenario in which we might expect atypical activation patterns) and the second examined the impact of removing trials with short Stop Signal Delays (SSD) from all participants. Short SSD trials (SSD < 150 msec), in which the Go stimuli were presented for the shortest durations, are likely to be the trials driving any context independence violations. The first analysis compared Stop activation (the contrasts Successful Stops and Successful Stops vs Correct Go trials) between the group of violators (n = 257; baseline [ages 9 and 10] data only) and non-violators (n = 5,001). Covariates included sex, age (in months), highest parental education, race/ethnicity, puberty level, and scanner. No group differences were observed with a vertex-wise threshold of p < .05 and application of either a family-wise error correction based on random field theory or a less conservative false discovery rate. At a nominal p < .05, uncorrected threshold, group differences were observed in visual cortex only. Further, in a region-of-interest analysis of the right IFG, a critical node of the response inhibition network, violators and non-violators did not differ in activation (p = .83).

To quantify similarity between groups in the spatial patterns of activation, we calculated the vertex-wise correlation between group activation of the violators and a “gold standard” of activation based on the remaining non-violator participants (the whole sample was first residualized for the covariates listed above). To facilitate interpretation, we quantified the similarities that would be expected with samples of size 257 by comparing randomly selected subsamples of non-violators (n = 257, 10,000 samples) against the remainder (n = 4,487; Figure 2A). Figure 2b shows the distribution of vertex-wise correlations for both the Successful Stops contrast (blue) and the Successful Stops vs Correct Go contrast (orange). The correlations for violators are indicated by the red line. Although lower than the mean of the subsampling distribution, shown by the dotted blue line, we note that the vertex-wise correlation is very high, even for violators (r = .94 and .94 for violators, compared to the full group of non-violators with r = .95 and .96, for Successful Stops and Successful Stops vs Correct Go, respectively). Moreover, violators, unsurprisingly, show relatively poor performance on the task (Figure 2c). An equal sized group of non-violators, matched to the violators on Stop success rate, Stop Fail accuracy, Go RT, Go accuracy, and Go omission rate are indicated by the green lines (Matched Group). We conclude that even those participants identified as violating race model assumptions show activation patterns that are very similar to those observed for non-violators.

**Figure 2:**
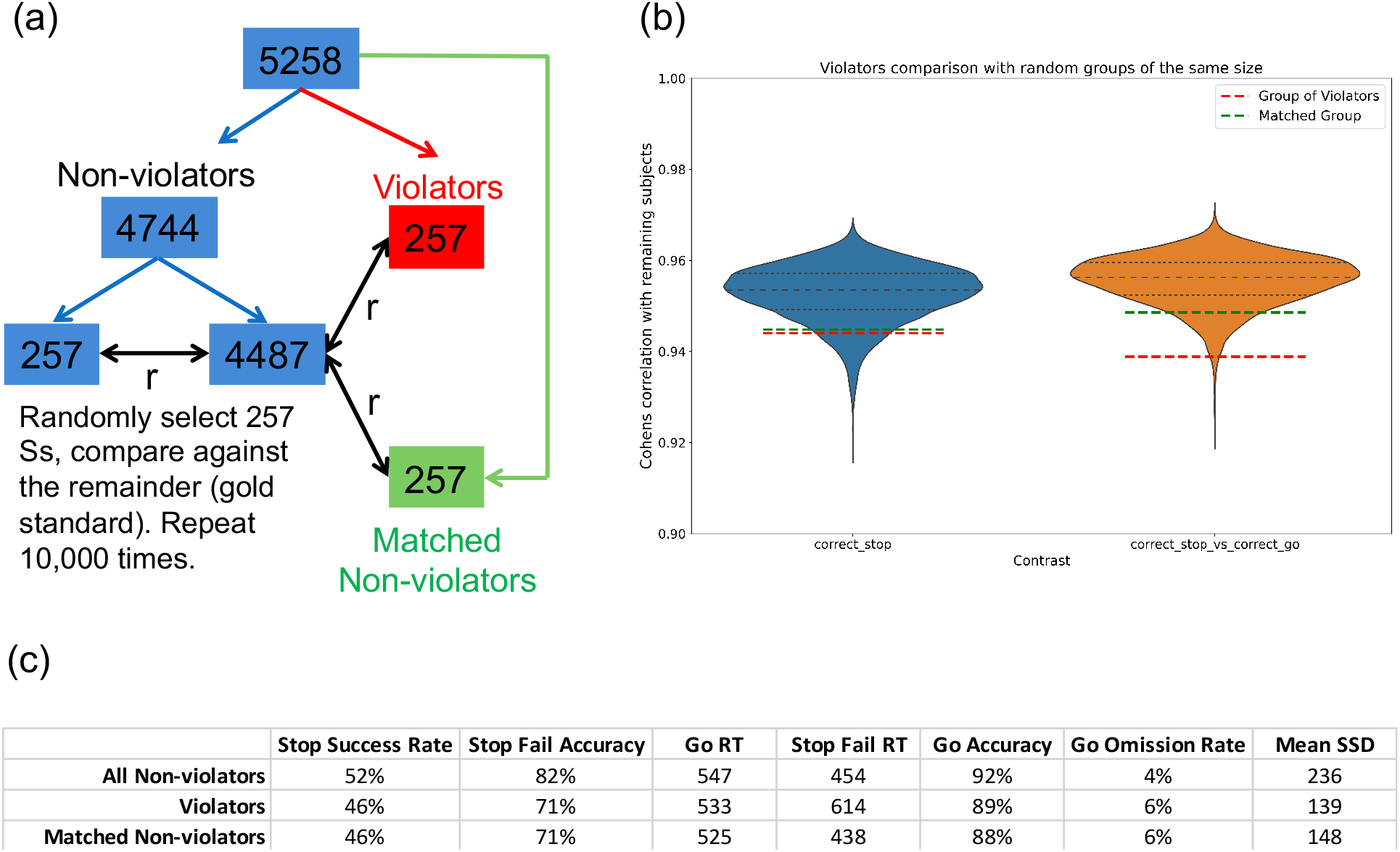
Correlation of violators and performance-matched non-violators with gold-standard brain activation. (a) A subsampling procedure determined the similarity between vertex-wise activation levels in samples of n=257. (b) Correlations with the “gold standard” activation map for subsamples of non-violators (blue and orange distributions), violators (red line) and performance-matched non-violators (green line). (c) Performance of violators, all non-violators, and performance-matched non-violators.

The second analysis compared group Stop activation maps (Successful Stops vs Correct Go) with all trials versus with the shorter SSD Stop trials (0msec, 50msec, 100msec) excluded. The same covariates as described above were included (n = 5,058). Although the amplitude of activation was larger in the former (due, presumably, to the inclusion of more trials), critically, the patterns of activation were almost identical (Figure 3). The vertex-wise correlation between the group activation map that included all trials with the group activation map that excluded the 0msec, 50msec, and 100msec SSD trials was r = .99.

**Figure 3:**
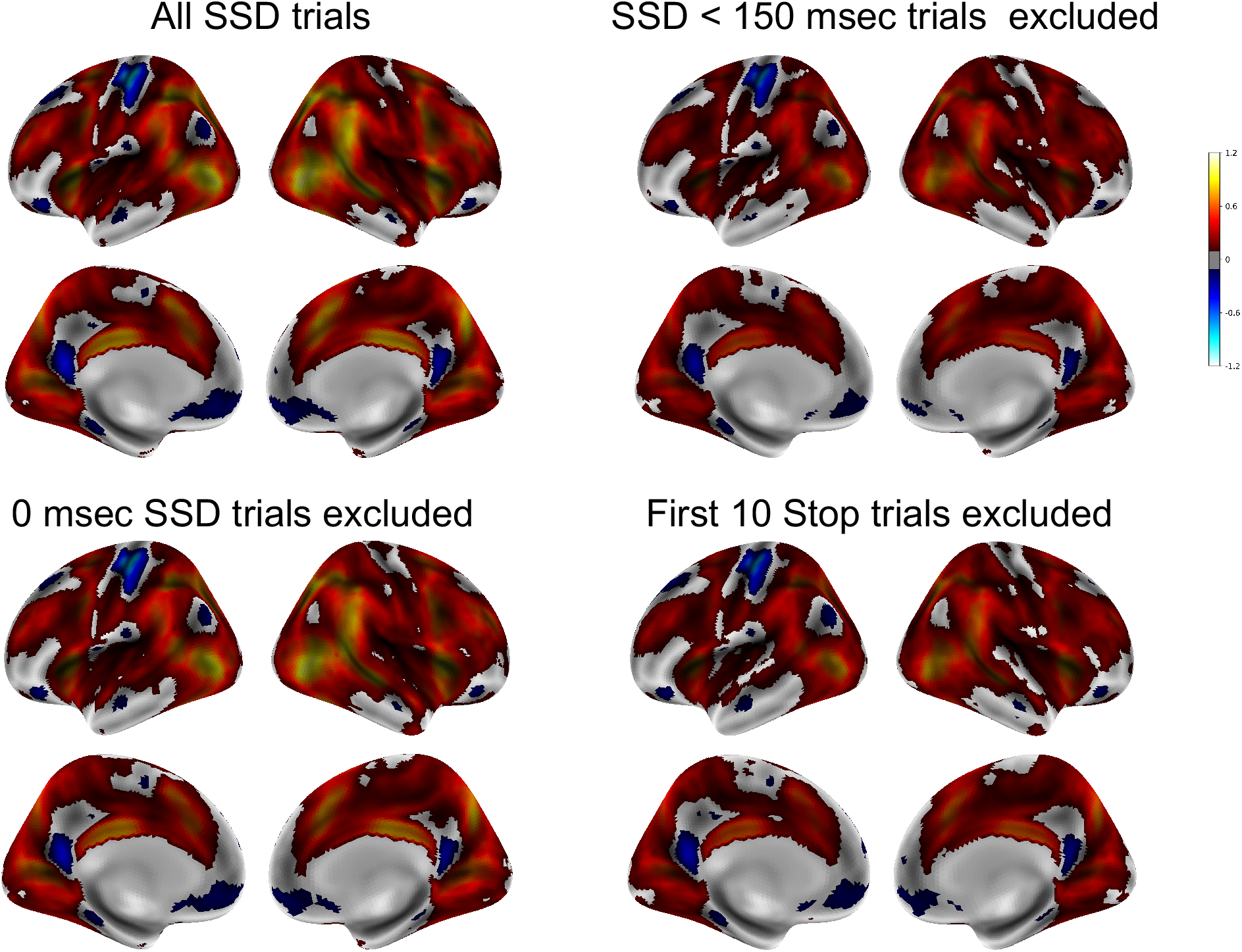
Brain activation (Cohen’s d threshold of .2 for Successful Stops vs Correct Go trials) for all Stop trials, Stop trials with SSD = 0/50/100 msec excluded, Stop trials with SSD = 0 msec excluded, and first ten Stop trials excluded.

#### Issue 1 Implications and Recommendations

Race model violators (Stop Fail RT > Go RT) are now identified in annual data releases and we suggest that researchers not include them when estimating SSRT (Verbruggen et al. 2019). We do not believe that there is sufficient evidence at this stage to warrant distrust of the remaining performance and neuroimaging data but encourage investigators to consider the impact of the issues that have been raised for their research question and their application of SSRT measurement models, and to apply what they deem to be appropriate inclusion and exclusion criteria. The ABCD task fMRI working group, in consultation with its External Scientific Board and additional external experts, have decided that changes to the fundamental task design are not warranted at this stage. The group notes that the poorer choice accuracy on Stop Fail trials does indicate a degree of independence violation that likely would be reduced if the Go stimuli remain on screen (for the 1 sec duration of the trial or until a response is made). However, noting that context independence violations are common across numerous task designs (Bissett et al., 2021), the analyses reported above suggest that the current task is yielding valuable, useful data that would appear to be largely unaffected by the concerns raised by Bissett and colleagues. Clearly, an ongoing longitudinal study will prioritize not changing a task without compelling, and ideally empirical, reasons to do so.

### Issue 2. Go stimulus sometimes not presented

Arising from the stimulus design feature underlying Issue 1, Go choice stimuli are not presented on trials in which the SSD drops to 0 msec. Bissett et al. suggest that this may confuse participants or “may make successfully stopping trivial (as the go process never started).” Bissett et al. report that 9.1% of Stop trials are 0 msec SSD trials but once the performance flagged participants and the Issue 3 programming error participants are excluded this drops to 6.9%. Curiously, performance on these trials is not trivial (average successful inhibition rate across participants on these trials is 64.8%) reflecting, presumably, the response prepotency induced by the high proportion of Go trials (see Table 2) which suggests that these trials also engaged inhibitory control processes.

**Table 2:** The percentage of Stop trials in which the SSD = 0msec and the probability of successfully inhibiting a response on these trials.

|  | All Ss | Exclude poor performers | Issue 3 exclusion |
| --- | --- | --- | --- |
| % of trials | 9.1% | 7.3% | 6.9% |
| Stop Success Rate | 60.3% | 63.7% | 64.8% |

The 0 msec SSD trials are broadly distributed across participants, with 49.9% of participants having at least one 0 msec SSD trials. We calculated SSRT with these 0 msec SSD trials included (283±78) and excluded (276±73). (The average of a participant’s SSDs is included in the calculation of their SSRT, so the exclusion of all 0 msec SSD trials would be expected to produce a shorter SSRT estimate.) Notably, the correlation between the two estimates is very high (r = .97). The broad distribution of these trials across participants appears to reduce their impact on subsequent analyses. Turning to the neuroimaging data, the vertex-wise correlation between brain activation when these trials are included vs. excluded is very high (r = .99; see Figure 3). This correlation holds when all participants are included (n = 5,064) and when the analyses are restricted to those participants with one or more 0 msec SSD Stop trials (n = 2,416).

#### Issue 2 Implications and Recommendations

To avoid potential confusion, the task has been modified to ensure that the SSD does not drop below 50 msec thereby ensuring presentation of the Go stimulus on all trials. The impact of the 0 SSD trials on the existing data appears to be very small. For researchers who may wish to exclude participants with a high number of these trials, we now include the number of 0 msec SSD trials per participant in annual data releases.

### Issue 3. Degenerate stop-signal delays

Bissett et al. identified a programming error and we thank them for bringing this to our attention. When the SSD is 50 msec, a response that is faster than 50 msec is erroneously recorded as the response for all subsequent Stop trials. Bissett et al. report that this programming error affects 2.67% of participants. However, if the performance flags are applied this reduces to 1.24% of participants. The data of many of these participants are likely retrievable if one restricts analyses to the data obtained prior to the onset of the error. For example, just 0.8% of participants have this problem occur prior to their 50^th^ Stop trial and the “best practices” paper by Verbruggen and colleagues concludes from a series of simulations that “reliable and unbiased SSRT group-level estimates can be obtained with 50 stop trials.”

#### Issue 3 Implications and Recommendations

The programming error in the task has been corrected and the corrected task is available on the ABCD study website (https://abcdstudy.org/families/abcd-fmri-tasks-and-tools/). For the existing data, although we anticipate that valuable performance and brain imaging data of many participants affected by this programming error are retrievable, we include a variable identifying these participants in data releases, enabling researchers to exclude them from analyses.

### Issue 4. Different Stop Signal duration for different SSDs

This issue arose because all trial events were constrained to happen within the 1 second trial period and not to carry into the inter-trial interval. As a consequence, the duration of the Stop signal (typically 300 msec) was shortened if the SSD was greater than 700 msec. One might expect these events to be quite uncommon and also to be indicative of other performance-related problems (i.e., a 700 msec SSD is atypically long). We calculate the frequency of these shorter Stop signal durations to be 1.15% of all Stop trials. However, this reduces to 0.12% of trials once performance criteria are applied (as mentioned, the presence of these very long SSD trials indicates other performance problems). Moreover, as participants very often respond during the relatively long SSD, the proportion of trials in which the shorter SSDs are presented and on which participants have not already responded reduces to 0.07% of trials. We removed these short Stop signal duration trials and assessed the impact on SSRT. Identical results were obtained: With all trials included and with these short Stop signal duration trials included, SSRT = 283±78.

#### Issue 4 Implications and Recommendations

Given the rarity of these trials, we conclude that they have a negligible impact on the SSRT estimates. Nonetheless, we have changed the task to ensure that the Stop signal duration is always 300 msec in duration, a change which we believe will have a negligible impact on any longitudinal comparisons.

### Issue 5. Non-uniform conditional trial probabilities

Bissett et al. note correctly that the trial orders (Stop vs. Go) were not fully randomized (in accordance with the ratio of Stop to Go trials). Instead, the conditional trial probabilities and inter-stimulus intervals (ISIs) for ABCD were selected to optimize the joint estimation efficiency for fMRI responses to Go and Stop events, assuming a canonical hemodynamic response function. For each of 40 runs, sequences of Stop and Go stimuli, and randomized jittered intervals between them, were selected by generating 100,000 random trial sequences and ISIs (constrained to 0.7 – 2 sec, starting mean 0.9 sec), and choosing the design with the minimum mean Variance Inflation Factor. The design was constrained to avoid repeating successive Stop trials. The rationale for this is based on evidence that repeated Stops are easier, thereby reducing power and homogeneity in the demand on inhibition across trials (Bissett et al., 2012).

Bissett et al. raise concerns that the non-uniform trial probabilities might be learned by participants. Their analysis of post-Stop RTs indicate that this is not the case (i.e., initial post-Stop slowing does not transition, as the task progresses, to post-Stop speeding up) but they caution that it may be learned as the participants age. A very similar Stop task (i.e., no repeated Stop trials) has been used in the IMAGEN longitudinal study of ~2,000 adolescents. We assessed post-Stop behavior similar to Bissett et al. (post-Stop RT minus pre-Stop RT, separating all Stop trials into quartiles) when these participants performed the task at age 19, which was the second administration of the task to them. Figure 4 shows a very similar pattern in IMAGEN to what Bissett et al. report for ABCD. There is no evidence that participants at age 19 learn to speed up on post-Stop trials and, instead, post-Stop slowing is maintained.

**Figure 4:**
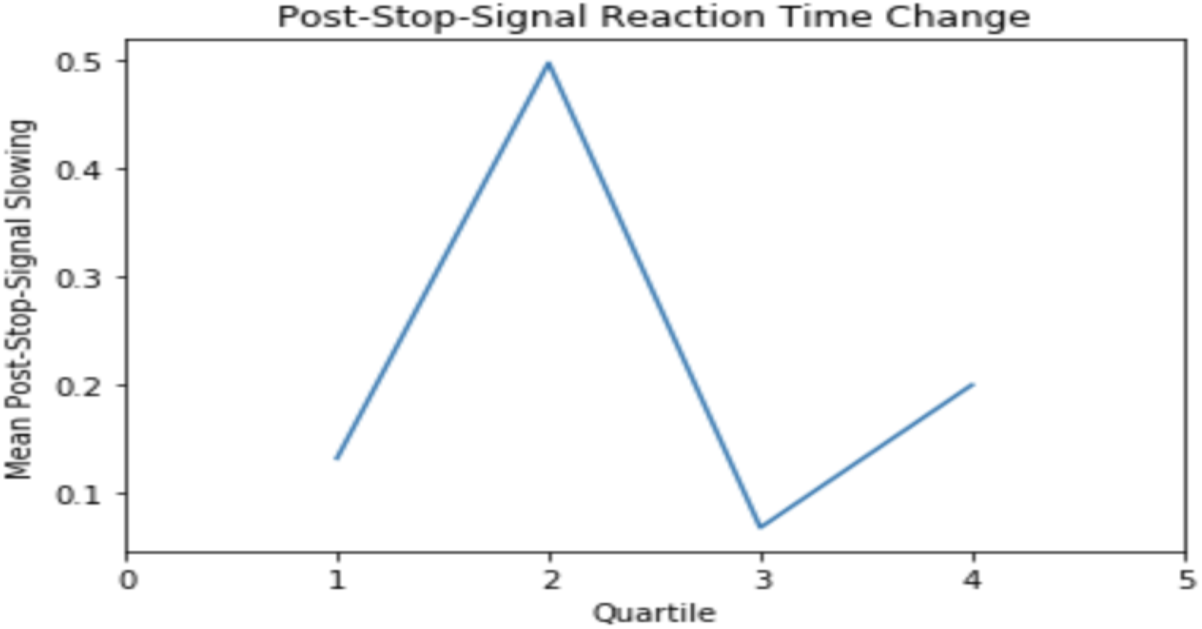
Post-Stop slowing (i.e., RT in seconds on the Go trial that immediately followed a Stop trial minus RT on the Go trial that immediately preceded the Stop trial) is shown across trial quartiles for the IMAGEN participants at age 19. This task also excludes repeated Stop trials and, similar to the younger ABCD participants, shows no evidence of changes in post-Stop slowing between the start and end of the task.

#### Issue 5 Implications and Recommendations

We recommend no changes to the task design. Researchers concerned about trial conditional probabilities being learned as the ABCD participants age can assess these in a manner as suggested by Bissett et al. Indeed, the extent to which participants develop subjective expectations and prepare for Stop trials is an interesting avenue of research for investigators interested in proactive control.

### Issue 6. Trial accuracy incorrectly coded

Bissett et al. describe a number of trial outcome labeling errors. Unfortunately, Bissett et al did not explain that all of these errors had already been identified by the ABCD Data Analysis and Informatics Resource Center, that they were corrected and documented prior to data analysis and data release, and that the code to make these corrections was shared publicly. These errors derived from how E-Prime pre-release settings were used: Responses occurring during the pre-release period of the post-trial jittered intervals were not recorded correctly, resulting in Stop Fail trials being logged as Stop Successes and Correct Go trials being logged as Go Omissions. Details on the trial labeling errors and their correction are included in the data processing scripts available from the ABCD github site (https://github.com/ABCD-STUDY/). The misclassification of errors on the Stop trials (classifying what should be Stop Fails as Stop successes) did impact the task (i.e., SSD increased when it should have decreased), but this occurred on only a small fraction of trials and, as Bissett et al. report, would have had a very small effect on the SSD tracking algorithm.

#### Issue 6 Implications and Recommendations

The cause of the errors has been corrected in the task code. Data available through the annual NDA releases have already corrected for these labelling errors but researchers wishing to work with the raw E-Prime output are encouraged to employ the corrections specified in the “abcd_extract_eprime_sst” script available on the ABCD github site.

### Issue 7. SSD values start too short

Bissett et al. query the reasoning behind starting the task with the SSD set to 50 msec rather than a value that is closer to what the final SSD would be (e.g., 250msec). The shorter SSD starts the task at a relatively easy level thereby providing a “warm-up” of sorts, easing participants, aged just 9 and 10 at baseline, into what is a cognitively challenging task. As shown in Figure 6 of Bissett et al., these first few trials are, in fact, not trivial for participants insofar as performance is better than average but far from ceiling (starting at 75% and dropping to 60% by the fifth Stop trial). Consequently, these first few trials contribute usefully to the behavioral and activation measures. As shown in Figure 1, the 60 Stop trials of the ABCD task are sufficient for the adaptive algorithm to converge on ~50% Stop success rate. To assess the impact of these starting trials, we removed the first ten Stop trials and all Go trials up to the tenth Stop trial and assessed the impact on SSRT. With all trials included SSRT = 283±78 and with the first ten Stop trials excluded SSRT = 284±85 with a high correlation between the two (r = .98). Similarly, group brain activation with and without these first ten Stop trials was calculated (n = 5,067) with the vertex-wise correlation between the two being very high, r = .99 (see Figure 3).

#### Issue 7 Implications and Recommendations

We believe there are no implications for the task or the data arising from this design feature.

### Issue 8. Low stop trial probability

The ratio of Go to Stop trials is 300:60 (17%) in ABCD, a ratio that increases task demands (i.e., increases the prepotency to respond, making inhibitions more difficult) which should ensure that this task continues to challenge participants as they age. We note that the task contains 60 Stop trials (as mentioned above, 50 are deemed sufficient for estimating SSRT) and successfully converges on ~50% Stop success rate (Figure 1). As noted above, a very similar task, with the same Stop trial probability has been used in the largest adolescent neuroimaging longitudinal study to date (IMAGEN; n = 2,000, with assessments at age 14, 19, and 23; Schumann et al., 2010) and it has demonstrated robust Stop success and Stop fail activations at ages 14 and 19 (age 23 data not yet available) and has demonstrated its ability to discriminate among adolescent phenotypes and predict future adolescent behavior (Whelan et al., 2012; 2014).

#### Issue 8 Implications and Recommendations

We believe there are no implications for the task or the data arising from this design feature. If researchers wish to compare the task’s activation levels or SSRT estimates against other datasets then, as is always the case in comparing separate studies, they should be aware of the particular design features of the ABCD study.

## Conclusion

We encourage researchers to continue analyzing the ABCD data and to help us improve the study by identifying potential problems like Bissett and colleagues have done. In light of the concerns raised by Bissett et al, some changes to the ABCD Stop task have been made (Table 3). Quantifying the impact of the specific concerns on the validity of the data is, of course, challenging: Although a specific cause for context violations can be identified in the ABCD Stop task, context violations as Bissett and colleagues have noted elsewhere (Bissett et al., 2021) are, in fact, quite widespread across multiple task designs and, consequently, researchers must always attend to this and other assumptions of their measurement models and analyses. The analyses presented here lead us to conclude that the specific design feature of the ABCD Stop task appears, thus far, to have a minimal impact on the neuroimaging data. The impact on the SSRT data, including on the rank ordering of participants, appears to be modest, especially if the recommendations provided here (which does include some new participant exclusions) are followed. That said, we await more empirical and computational analyses on these matters and encourage researchers to consider the implications of the task design for the analyses they conduct and any measurement model they apply to these data. More generally, we encourage researchers to contact the ABCD team promptly should their analyses raise concerns with any element of the assessment battery. Doing so ensures that misunderstandings can be avoided and any errors speedily corrected.

**Table 3:**
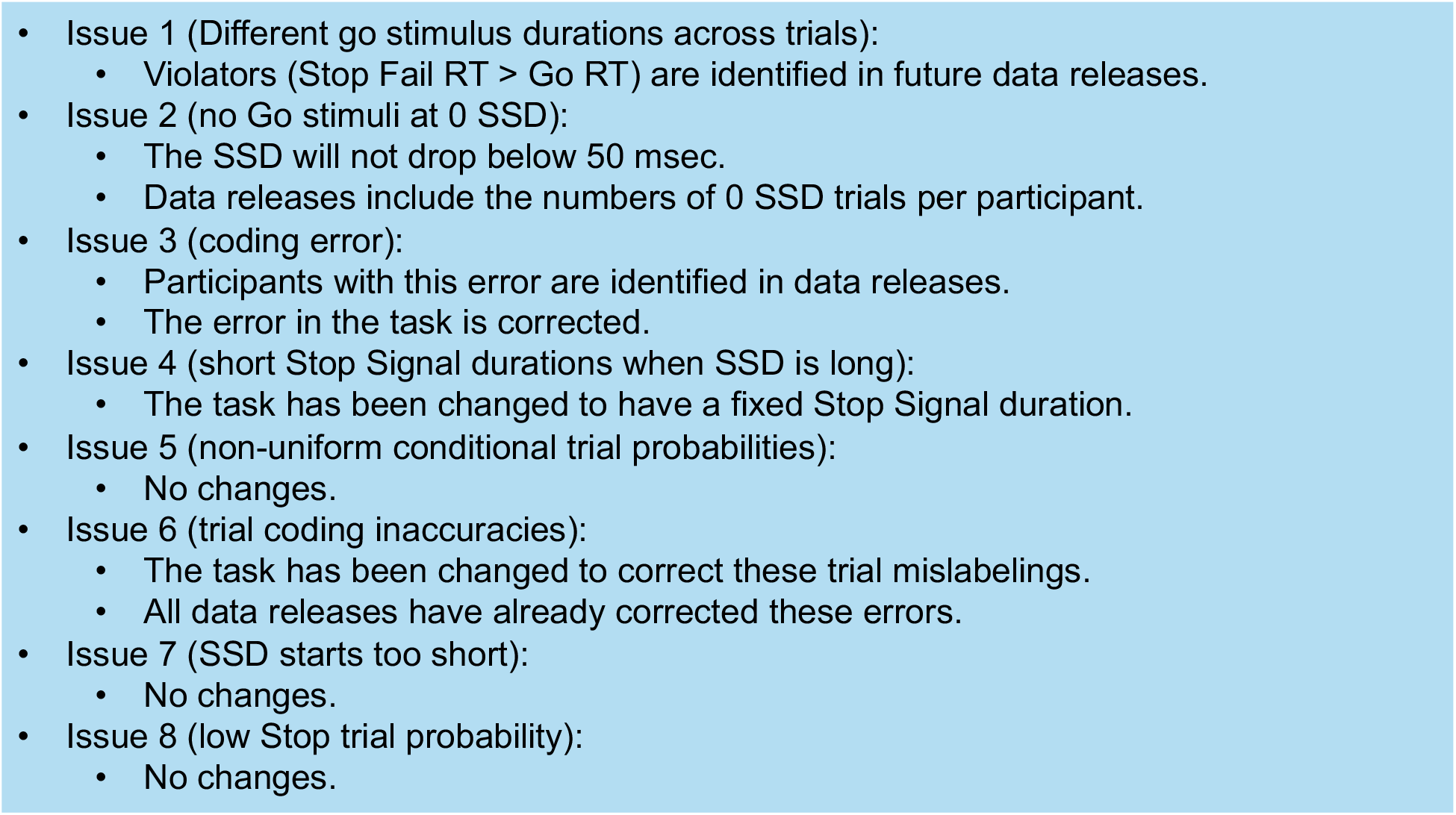
Task and data sharing changes.

## Acknowledgements

We thank Harriet de Wit, Ted Satterthwaite, Alex Weigard, members of the ABCD External Scientific Board, and members of the ABCD task fMRI working group for their helpful discussions of these topics.

## Footnotes

1 Excluding participants who fail to adequately comply with task instructions or who show aberrant performance is a standard practice and especially important given the young age of ABCD participants. Recommended performance criteria that accompany ABCD data releases are shown in Figure 1 but these were not applied by Bissett et al. In addition, a programming error detailed under Issue 3 led to a very different task experience for certain participants and we are recommending that these participants (1.24% of the sample) be excluded from analyses.

